# Experimental signal dissection and method sensitivity analyses reaffirm the potential of fossils and morphology in the resolution of the relationship of angiosperms and Gnetales

**DOI:** 10.1101/134262

**Authors:** Mario Coiro, Guillaume Chomicki, James A. Doyle

## Abstract

The placement of angiosperms and Gnetales in seed plant phylogeny remains one of the most enigmatic problems in plant evolution, with morphological analyses (which have usually included fossils) and molecular analyses pointing to very distinct topologies. Almost all morphology-based phylogenies group angiosperms with Gnetales and certain extinct seed plant lineages, while most molecular phylogenies link Gnetales with conifers. In this study, we investigate the phylogenetic signal present in published seed plant morphological datasets. We use parsimony, Bayesian inference, and maximum likelihood approaches, combined with a number of experiments with the data, to address the morphological-molecular conflict. First, we ask whether the lack of association of Gnetales with conifers in morphological analyses is due to an absence of signal or to the presence of competing signals, and second, we compare the performance of parsimony and model based approaches with morphological datasets. Our results imply that the grouping of Gnetales and angiosperms is largely the result of long branch attraction, consistent across a range of methodological approaches. Thus, there is a signal for the grouping of Gnetales with conifers in morphological matrices, but it was swamped by convergence between angiosperms and Gnetales, both situated on long branches. However, this effect becomes weaker in more recent analyses, as a result of addition and critical reassessment of characters. Even when a clade including angiosperms and Gnetales is still weakly supported by parsimony, model-based approaches favor a clade of Gnetales and conifers, presumably because they are more resistant to long branch attraction. Inclusion of fossil taxa weakens rather than strengthens support for a relationship of angiosperms and Gnetales. Our analyses finally reconcile morphology with molecules in favoring a relationship of Gnetales to conifers, and show that morphology may therefore be useful in reconstructing other aspects of the phylogenetic history of the seed plants.

## INTRODUCTION

The use of morphology as a source of data for reconstructing phylogenetic relationships has lost most of its ground since the advent of molecular phylogenetics, except in paleontology. However, there has recently been renewed interest in morphological phylogenetics (Pyron 2015; Lee and Palci 2015). This is partly because of increased focus on the phylogenetic placement of fossil taxa in trees of living organisms, stimulated by the necessity of accurate calibrations for dating the molecular trees that have become the main basis for comparative evolutionary studies. This has led to the development of methods that integrate phylogenetic placement of fossils in the dating process (Pyron 2011; Ronquist et al. 2012; Zhang et al. 2016). Another focus has been the application of statistical phylogenetics to morphological data on both a theoretical (Wright et al. 2014, 2015; O’Reilly et al. 2016) and an empirical level (Lee and Worthy 2012; Godefroit et al. 2013; Cau et al. 2015). In paleontology, where only morphological data are available (except in the recent past), questions on the role of morphology in phylogenetics are even more critical. A major issue concerns the value of fossils in reconstructing relationships among living organisms. Early in the history of phylogenetics, there were claims that fossils are incapable of overturning phylogenetic relationships inferred from living taxa (Patterson 1981), but also demonstrations that they can, as for instance in morphological analysis of amniote phylogeny (Gauthier et al. 1988). Whether or not fossils affect the inferred topology of living taxa, there is little doubt that they are often either useful or necessary in elucidating the homologies of novel structures (e.g., the seed plant ovule and eustele) and the order of origin of the morphological synapomorphies of extant (crown) groups (e.g., origin of secondary growth before the ovule in the seed plant line), as discussed in Doyle (2013). This is critical because major groups, such as the now-dominant angiosperms (flowering plants), are often separated from their closest living relatives by major morphological gaps (numbers of character changes), even if the incorporation of fossils does not affect inferred relationships among living taxa (Doyle and Donoghue 1987; Donoghue et al. 1989).

Many phylogenies based on morphology have been recently published for important groups with both living and fossil representatives, including mammals (O’Leary et al. 2013), squamate reptiles (Gauthier et al. 2012), arthropods (Legg et al. 2013), and the genus *Homo* (Dembo et al. 2016). However, the validity and use of morphological data in reconstructing phylogeny have been severely criticized, notably by Scotland et al. (2003), based on supposed diminishing returns in the discovery of new morphological characters and the prevalence of functional convergence. The painstaking acquisition of morphological characters, which requires a relatively large amount of training and time, could turn out to be systematically worthless if the phylogenetic signal present in these data is either insufficient or misleading. Indeed, the number of characters that can be coded for morphological datasets represents a major limit to the use of morphology and its integration with molecular data, especially in the age of phylogenomics, where the ever-increasing amount of molecular signal could simply “swamp” the weak signal present in morphological datasets (Doyle and Endress 2000; Bateman et al. 2006). Morphological data may also be afflicted to a higher degree than molecules by functional convergence and parallelism (Givnish and Sytsma 1997), which could lead a morphological dataset to infer a wrong phylogenetic tree. Even though the confounding effect of convergence has been formally tested in only a few studies (Wiens et al. 2003), it seems to be at the base of one of the deepest cases of conflict between molecules and morphology in the reconstruction of evolutionary history, namely the phylogeny of placental mammals (Foley et al. 2016). In this case, the strong effect of selection on general morphology caused by similar lifestyle seems to hinder attempts to use morphology to reconstruct phylogenetic history in this group (Springer et al. 2007), and it affects even large “phenomic” datasets (Springer et al. 2013).

Another example of conflict between morphology and molecular data involves the relationships among seed plants, particularly angiosperms and the highly derived living seed plant order Gnetales. Before the advent of cladistics, some authors proposed that angiosperms and Gnetales were closest living relatives, while others argued that these two groups were strictly convergent and Gnetales were instead related to conifers (for a review, see Doyle and Donoghue 1986). However, since the earliest studies by Parenti (1980) and Hill and Crane (1982), which included only living taxa, the view that angiosperms are most closely related to Gnetales has appeared to be one of most stable results of morphologically based parsimony analyses of seed plant phylogeny (Crane 1985a; Doyle and Donoghue 1986, 1992; Nixon et al. 1994; Rothwell and Serbet 1994; Doyle 1996, 2006, 2008; Hilton and Bateman 2006; Friis et al. 2007; Rothwell et al. 2009; Rothwell and Stockey 2016; Fig. 1). The first analysis that included fossils, by Crane (1985a), associated angiosperms and Gnetales with Mesozoic Bennettitales and *Pentoxylon*. Because all four taxa have more or less flower-like reproductive structures, this clade became known as the anthophytes, a term formerly used for angiosperms, to emphasize its implication that the flower was a synapomorphy not of angiosperms alone but rather of a larger clade to which they belong (Crane 1985b; Doyle and Donoghue 1986). Some subsequent analyses interpolated the Mesozoic fossil *Caytonia* into this clade as the closest outgroup of angiosperms (Doyle 1996, 2006, 2008; Hilton and Bateman 2006; Friis et al. 2007). This result calls into question the original concept of anthophytes as a clade united by flowers, since *Caytonia* had large sporophylls that are unlikely to have been grouped into flower-like structures. However, all trees found in morphological analyses, with the exception of some in Doyle (2008), have agreed that Gnetales are the closest living relatives of angiosperms. Some analyses associated the clade including angiosperms and Gnetales with “Mesozoic seed ferns” (such as glossopterids, corystosperms, and *Caytonia*), others with “coniferophytes” (conifers, *Ginkgo*, and fossil cordaites). Inferred relationships within the clade have also varied: in some cases angiosperms and Gnetales are sister groups, in others Gnetales are linked with Bennettitales. Because some studies place taxa without flower-like structures in the clade, and molecular data are unable to distinguish such trees from anthophyte trees in the original sense, we refer to the whole class of trees in which angiosperms and Gnetales are closest living relatives as “gnetangiosperm” rather than “anthophyte” trees.

**Figure 1.**
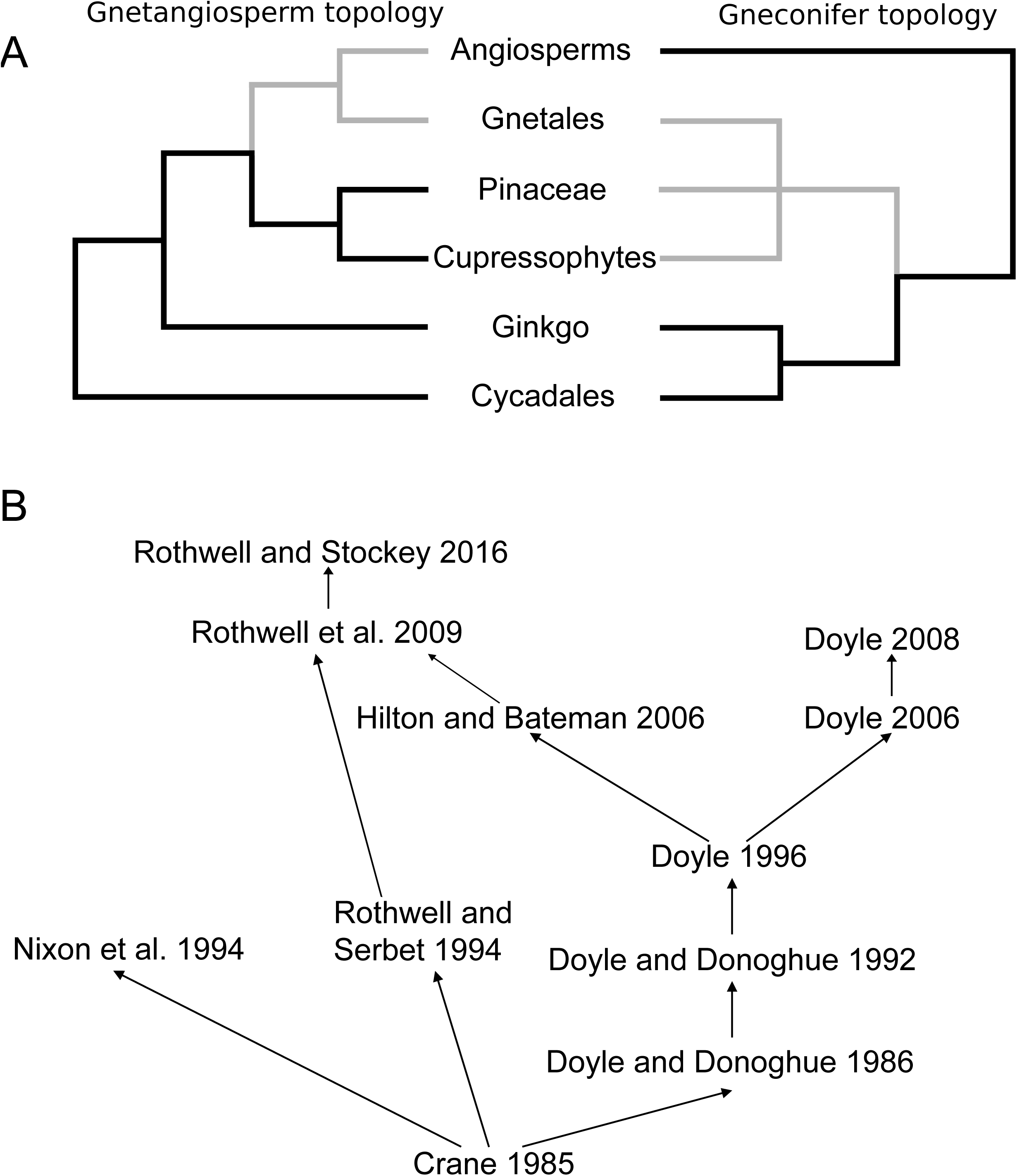
A, Relationships among extant seed plants. On the left an gnetangiosperm topology, and on the right a gneconifer topology. Relationships between Cycadales and *Ginkgo* vary among analyses of both sorts. B, Relationships among the matrices reanalyzed in this paper.

By contrast, since the advent of molecular phylogenetics, the hypothesis that angiosperms and Gnetales are closely related has lost most of its support among plant biologists. Although molecular analyses cannot directly evaluate the status of putatively related fossil taxa, they can address the relationship of angiosperms and Gnetales. Molecular data from different genomes analyzed with different approaches do not yield a Gnetales plus angiosperm clade, with the exception of few maximum parsimony (MP) and neighbor joining analyses of nuclear ribosomal RNA or DNA (Hamby and Zimmer 1992; Stefanovic et al. 1998; Rydin et al. 2002) and one MP analysis of *rbcL* (Rydin and Källersjö 2002). The majority of molecular analyses retrieve a clade of Gnetales plus Pinaceae (Bowe et al. 2000; Chaw et al. 2000; Gugerli et al. 2001; Qiu et al. 2007; Zhong et al. 2011), conifers other than Pinaceae (cupressophytes) (Nickrent et al. 2000; Rydin and Källersjö 2002), or conifers as a whole (Wickett et al. 2014), which we refer to collectively as “gneconifer” trees. In most of these trees angiosperms are the sister group of all other living seed plants (acrogymnosperms: Cantino et al. 2007). The main exceptions are “Gnetales-basal” trees, in which Gnetales are sister to all other living seed plants (e.g., Albert et al. 1994; Rydin and Källersjö 2002).

Several potential issues have been identified with both sorts of data. Regarding molecules, these include limited taxonomic sampling resulting from extinction of the majority of seed plant lineages (Rothwell et al. 2009), loss of phylogenetic signal due to saturation (particularly at third codon positions), strong rate heterogeneity among sites across lineages and conflict between gene trees (Mathews 2009), composition biases among synonymous substitutions (Cox et al. 2014), as well as systematic errors and biases (Sanderson et al. 2000; Magallón and Sanderson 2002; Burleigh and Mathews 2007; Zhong et al. 2011), leading to a plethora of conflicting signals. In analyzing datasets that yielded Gnetales-basal trees, studies that have attempted to correct for these biases have generally favored trees in which Gnetales are associated with conifers (Sanderson et al. 2000; Magallón and Sanderson 2002; Burleigh and Mathews 2007). Regarding morphology, in addition to far more complex problems in definition of characters and the role of functional convergence in confounding relationships, it has been shown that different taxon sampling strategies (which can also cause problems in molecular studies: Rydin and Källersjö 2002), such as choice of the closest progymnosperm outgroup of seed plants (Hilton and Bateman 2006), can lead to different results concerning the rooting of the seed plants.

The conflict between molecules and morphology has led to different attitudes toward morphological data within the botanical community (Donoghue and Doyle 2000; Scotland et al. 2003; Bateman et al. 2006; Rothwell et al. 2009). Following suggestions of Donoghue and Doyle (2000), Doyle (2006, 2008) reconsidered several supposed homologies between angiosperms and Gnetales in the light of the molecular results. These studies and the analysis of Hilton and Bateman (2006) also incorporated newly recognized similarities between Gnetales and conifers, for example in wood anatomy (Carlquist 1996), as well as new evidence on the morphology of the seed-bearing cupules in fossil taxa. Other changes involved redefinition of characters to reduce potential biases. For example, when building a morphological matrix, dissecting a character into more character states may represent an improvement by distinguishing convergent states and avoiding bias toward particular phylogenetic hypotheses during primary homology assessment (Jenner 2004; Zou and Zhang 2016), although it may be disadvantageous because it leads to a lack of resolution when the number of states becomes excessive. In seed plants, there are many special factors that complicate character coding. Among living taxa, the assessment of homology is complicated by the plastic and modular nature of plant development (Mathews and Kramer 2012). Among fossil taxa, the mode of preservation of many key fossils has critical consequences for the amount of data available. This affects not only the number of missing characters, but also the process of primary homology assessment and character coding. Although these issues with coding are most severe in fossils preserved as compressions, such as *Caytonia* (Doyle 2008; Rothwell et al. 2009) and *Archaefructus* (Sun et al. 2002; Friis et al. 2003; Doyle 2008; Rudall and Bateman 2008; Endress and Doyle 2009; Doyle and Endress 2014), even fossil groups that are exquisitely preserved as permineralizations (e.g., Bennettitales) are not immune to conflicting interpretations (Friis et al. 2007; Rothwell et al. 2009; Crepet and Stevenson 2010; Doyle 2012, supplemental material; Rothwell and Stockey 2013; Pott 2016).

Despite careful reconsideration of potentially convergent traits between Gnetales and angiosperms, the conflict between morphological and molecular data appeared to persist, with most morphological parsimony analyses continuing to favor the gnetangiosperm hypothesis (Doyle 2006; Hilton and Bateman 2006; Rothwell et al. 2009). The possibility that morphological data are inadequate to resolve such a key aspect of the phylogeny of seed plants would represent a severe hindrance in understanding plant evolution, especially in the light of the small number of extant lineages that survived extinction during the Paleozoic and Mesozoic (Mathews 2009) and the great morphological gaps among the surviving lineages. However, there have been signs that the conflicts with molecular data are weakening: in the analysis of Doyle (2006), trees in which Gnetales were nested in conifers were only one step less parsimonious than gnetangiosperm trees, and in Doyle (2008) trees of the two types became equally parsimonious.

In this study, we attempt to elucidate the phylogenetic signal present in published morphological datasets of the seed plants, concentrating on the relationship of angiosperms and Gnetales. This is not the only aspect of seed plant phylogeny that varies among and between morphological and molecular analyses. Another case is whether ginkgophytes (now reduced to *Ginkgo biloba*) are related to conifers and cordaites, as part of a coniferophyte clade, or to cycads, as found in some molecular analyses. However, the question of angiosperms and Gnetales is probably of the broadest evolutionary interest and is especially likely to illustrate the general problem of long branch effects in highly derived groups. We first test whether the possibility of convergence between angiosperms and Gnetales represents a major problem by reanalyzing the matrices that incorporated earlier homology assumptions concerning characters of the two groups (i.e., the matrices compiled before the incoming of molecular results) and later matrices that revised such assumptions (the matrices of Doyle 2006 and Hilton and Bateman 2006, and datasets derived from them) and testing whether the signal and the relative support for the gnetangiosperm and gneconifer clades changed between these two sets of matrices. After revealing a more coherent signal supporting a gneconifer clade in the more recent matrices, we investigate whether the retrieval of a gnetangiosperm topology by parsimony analyses was at least partly due to methodological biases that could be overcome by using model-based methods. Hopefully these approaches may be useful in resolving cases of conflict between morphological and molecular data in other taxa, particularly those with significant fossil representatives.

## MATERIALS AND METHODS

### Matrices

The matrices of Crane (1985a, version two, in which Bennettitales and *Pentoxylon* were scored as having cupules potentially homologous with those of Mesozoic seed ferns), Doyle and Donoghue (1986, 1992), Nixon et al. (1994), Rothwell and Serbet (1994), and Doyle (1996, 2006, 2008) were manually coded from the respective articles. The Hilton and Bateman (2006) matrix was kindly provided by Richard Bateman. The matrices from Analysis 3 of Rothwell et al. (2009) and from Rothwell and Stockey (2016) were downloaded from the supplementary materials of the respective articles.

### Parsimony analyses

We performed parsimony analyses of all matrices with PAUP 4.0a136 (Swofford 2003), using the heuristic search algorithm with random addition of taxa and 1000 replicates. Bootstrap analyses were conducted using 10,000 replicates, using the “asis” addition option and keeping one tree per replicate (Müller 2005).

We also conducted analyses with a topological backbone constraint, forcing the Gnetales into a clade with the extant conifers and leaving the position of other living taxa and fossils unconstrained. Significant differences between the constrained and unconstrained topologies were evaluated using the Templeton test (Templeton 1983) as implemented in PAUP v. 4.0a136 (Swofford 2003). We investigated the effects of recoding characters by Doyle (2006, 2008) in more detail by using MacClade (Maddison and Maddison 2003) to compare the number of steps in each character on trees with Gnetales associated with angiosperms and associated with conifers.

### Model-based analyses

Our model-based analyses were all conducted using the Markov k-states (Mk) model (Lewis 2001). This model assumes that characters are in one of *k* states, are all independent of each other, and change stochastically along branches with equal rates for all possible transitions, with all changes being independent of each other (as a Markov process). Some of these assumptions have been criticized for being unrealistic when applied to morphological change (Lewis 2001; Wright et al. 2014). For example, the model is fully symmetrical; i.e., the probability of change from 0 to 1 is equal to the probability of change from 1 to 0, an assumption that is violated by Dollo characters (i.e., losses of complex structures that are unlikely to be regained). Even though some of these assumptions can be theoretically relaxed, and implementations of these relaxed models already exist in a Bayesian framework (Wright et al. 2014), we used the standard version of the model to simplify the analyses and allow a closer comparison with the maximum likelihood implementation.

### Maximum likelihood (ML)

Maximum likelihood analyses were conducted using RaxML v. 8.2.10 (Stamatakis 2014). Matrices were modified by recoding all ambiguities (e.g., 0/1 in a three-state character) as missing data, since the method cannot cope with ambiguous characters. Topology is inferred using branch lengths, which are estimated as the expected number of state changes per character on that particular branch. We conducted 1000 bootstrap replicates with a gamma-distributed rate variation, which models different rates across characters by employing a multiplier drawn from a discretized gamma distribution.

### Bayesian inference (BI)

Bayesian analyses relied on MrBayes v. 3.2.3 (Ronquist et al. 2012), under the Mk model. For each matrix, we conducted two analyses, one with an equal rate of evolution among characters and another with gamma-distributed rate variation. In both cases, we used the MK_pr-inf_ correction for parsimony informative characters. The analyses were run for 5,000,000 generations, sampling every 1000^th^ generation. The first 10,000 runs were discarded as burn-in. Posterior traces were inspected using Tracer (Rambaut and Drummond 2007).

### Model testing and rate variation

We also conducted stepping stone analyses (Xie et al. 2011; Ronquist et al. 2012) in order to evaluate the most appropriate model of rate variation among characters (equal rates vs. gamma distributed rates). These analyses allow us to estimate the marginal likelihood for different models with better accuracy than other measures (e.g., harmonic mean estimator). We used 4 independent runs with 2 chains with the default MrBayes parameters, run for 5,000,000 generations and sampling every 1000^th^ generation. Using the marginal likelihoods from the stepping stone analysis, we then calculated the support for the two models using Bayes factors (BF) (Kass and Raftery 1995).

### Exploring conflict in the data

To explore phylogenetic conflict in the data, we employed the software SplitsTree 4 (Huson and Bryant 2006). We used this program to visualize conflicts among the bootstrap replicates from the MP and ML analysis and among the posterior tree samples found with Bayesian inference. The software summarizes the sets of trees using split networks, which allow us to visualize all possible conflicting hypotheses. These diagrams should not be confused with networks derived from distance-based neighbor-joining analyses. A consensus network (Holland et al. 2004) was built using the “count” option, with the cut-off for visualizing the splits set at 0.05.

### Long branch attraction tests

We modified the matrices to perform tests for long branch attraction (LBA), following the suggestions of Bergsten (2005). Two matrices were created to test the potentially destabilizing effect of the two long-branched groups suspected to create this artifact, angiosperms and Gnetales, by alternately removing each of them (long branch extraction analysis, LBE). If the association of angiosperms and Gnetales is indeed a result of LBA, then the removal of one of them should significantly alter the placement of the other. To test further the hypothesis of an LBA artifact exerted by angiosperms, we followed a similar approach to the sampling experiment in Rota Stabelli et al. (2010): another matrix was created to elongate the branch subtending angiosperms by removing the three fossil taxa most commonly identified as angiosperm outgroups (*Pentoxylon*, Bennettitales, and *Caytonia*) (branch elongation analysis, BE). In the presence of a long branch attraction artifact, the support for the node including the two long branches (angiosperms and Gnetales) should increase with such an “elongation” of one of the two branches. To test the effect of including fossil data in the matrices, we created a set of matrices in which all fossil taxa were removed (extant experiment, EX). Because this should lead to elongation of the branches subtending the living groups, this situation should result in the worst possible condition for long branch artifacts, and thus lead to the strongest apparent support for the node including the two long branches.

### Morphospace analysis

To visualize morphological patterns in the different matrices, we conducted principal coordinates (PCO) analyses. We employed the maximum observed rescaled distance between all pairs of taxa to generate the ordination as obtained using the MorphDistMatrix function of the R package Claddis (Lloyd 2016). PCO analysis was conducted using the ‘cmdscale’ function from the stats package (R Core Team 2017). The taxa were then plotted on the first two PCO axes.

### Data availability

All data are available on figshare: https://figshare.com/s/9a4fc5d4accff8e62084.

## RESULTS

Our re-analyses of the historical morphological matrices of seed plants with parsimony resulted in trees identical to the published trees (Table 1). The MP trees and the consensus trees always show a gnetangiosperm clade (with or without *Caytonia*), with the exception of trees based on the Doyle (2008) matrix, in which gnetangiosperm and gneconifer topologies are equally parsimonious. Constraining Gnetales and conifers to form a clade always results in trees longer than the most parsimonious trees, except with the Doyle (2008) matrix (Table 2). The Templeton test of the best trees against the worst constrained trees (i.e., the most parsimonious constrained tree that is statistically most different from the most parsimonious unconstrained tree) does however show that this difference is only significant at the 0.05 level with the Nixon et al. (1994) matrix.

**1.** Statistics for the parsimony analyses of fossil matrices.

|  | *Number of*<br>*trees* | *Length* | *Ci* | *Ri* |
| --- | --- | --- | --- | --- |
| **Crane 1985** | 8 | 50 | 0.600 | 0.730 |
| **Doyle and Donoghue 1986** | 36 | 123 | 0.504 | 0.674 |
| **Doyle and Donoghue 1992** | 94 | 112 | 0.545 | 0.658 |
| **Nixon et al. 1994** | 225 | 332 | 0.392 | 0.788 |
| **Rothwell and Serbet 1994** | 8 | 191 | 0.529 | 0.721 |
| **Doyle 1996** | 123 | 247 | 0.494 | 0.782 |
| **Hilton and Bateman 2006** | 480 | 313 | 0.457 | 0.801 |
| **Doyle 2006** | 8 | 321 | 0.514 | 0.753 |
| **Doyle 2008** | 16 | 346 | 0.503 | 0.744 |
| **Rothwell et al. 2009** | 66 | 330 | 0.503 | 0.776 |
| **Rothwell and Stockey**<br>**2016** | 6 | 363 | 0.466 | 0.754 |

**Table 2.** Results from the MP analysis of constrained gneconifer trees

|  | *Length*<br>*uncons*<br>*trained* | *Length*<br>*Gnetales+Conif*<br>*er* | *Length*<br>*differenc*<br>*e* | *Templeton Test p-value*<br>*(best value)* |
| --- | --- | --- | --- | --- |
| **Crane 1985** | 50 | 54 | 4 | 0.1573 |
| **Doyle and**<br>**Donoghue**<br>**1986** | 123 | 130 | 7 | 0.1266 |
| **Doyle and**<br>**Donoghue**<br>**1992** | 112 | 118 | 6 | 0.1088 |
| **Nixon et al.**<br>**1994** | 332 | 348 | 16 | 0.0131* |
| **Rothwell and**<br>**Serbet 1994** | 191 | 197 | 6 | 0.2252 |
| **Doyle 1996** | 247 | 257 | 10 | 0.0679 |
| **Hilton and**<br>**Bateman 2006** | 313 | 317 | 4 | 0.4595 |
| **Doyle 2006** | 321 | 322 | 1 | 0.8474 |
| **Doyle 2008** | 346 | 346 | 0 | 0.9888 |
| **Rothwell et al.**<br>**2009** | 330 | 334 | 4 | 0.3458 |
| **Rothwell and**<br>**Stockey 2016** | 363 | 369 | 6 | 0.1336 |

Bootstrap analysis shows that the gnetangiosperm clade is not strongly supported by any of the matrices, with the exception of the Nixon et al. (1994) matrix (Fig. 2). In the MP bootstrap analysis of the post-2000 matrices (Fig. 2A), support for a gnetangiosperm topology appears to be lower than support for a gneconifer topology in all matrices except that of Rothwell et al. (2009). The ML bootstrap (Fig. 2B) shows higher support for a gneconifer topology than the MP bootstrap in all post-2000 analyses, as well as in the two pre-2000 Doyle and Donoghue (1986, 1992) matrices. In the post-2000 matrices, the support for gneconifers is always higher than the support for gnetangiosperms.

**Figure 2.**
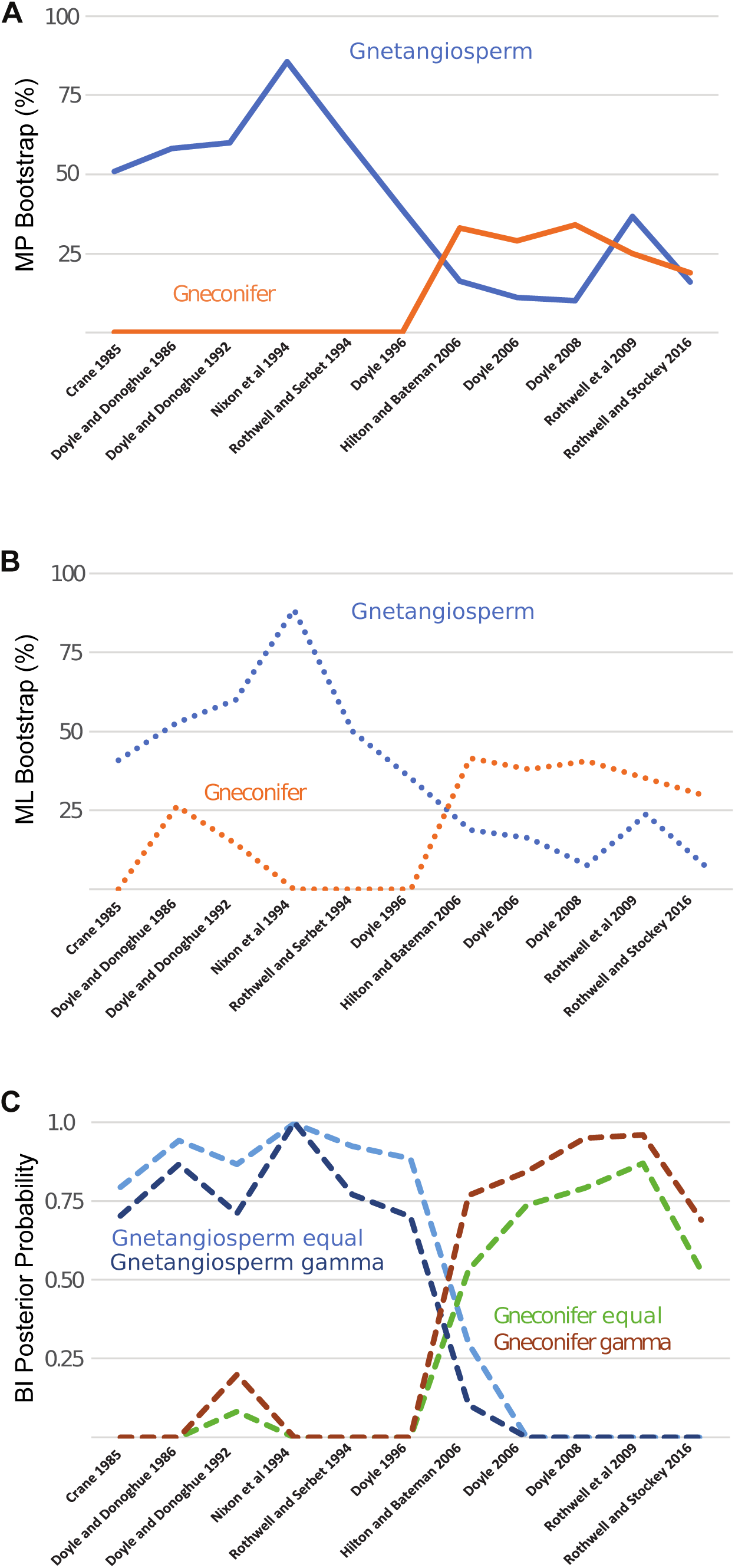
Support for the gnetangiosperms or gneconifers in the different matrices and using different methods. A, Results from the MP bootstrap analyses; B, results from the ML bootstrap analyses; C, results from the BI analyses. The difference between the pre-2000 and post-2000 matrices is clearly underlined by a shift in support from gnetangiosperms to gneconifers in the ML and BI analyses, and a drop in support for the gnetangiosperms in the MP analyses.

Our Bayes factor analysis using the marginal likelihood from the stepping stone runs shows strong support for rate variation among characters in all matrices except those of Crane (1985a) and Doyle and Donoghue (1986) (Table 3), as indicated by ln-Bayes factors higher than 2.

**Table 3.**
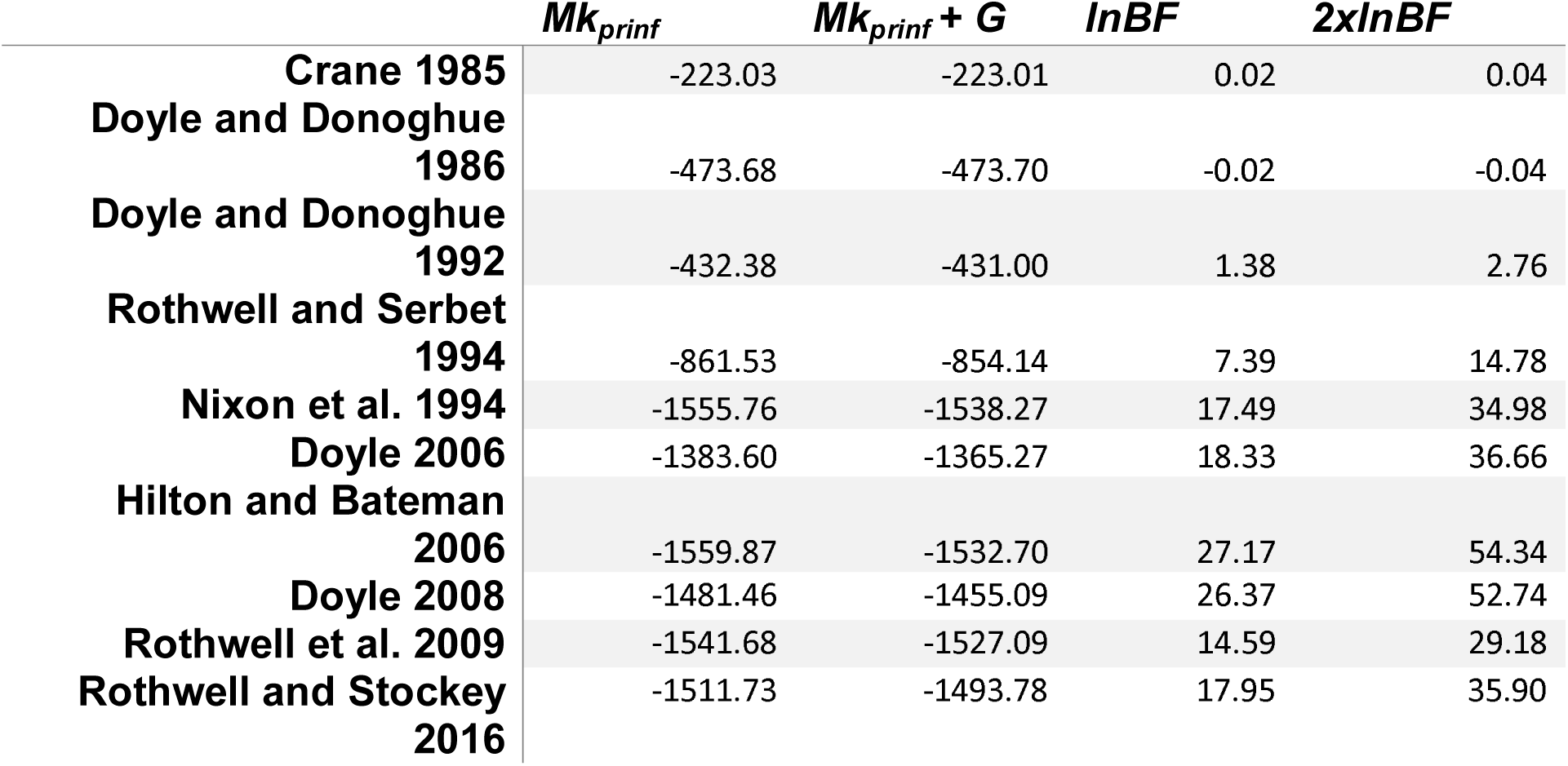
Model-testing statistics for the Bayesian inference analyses.

The trees obtained from the Bayesian analyses show a much sharper differentiation between early and late matrices, as shown by the trends in support values for gnetangiosperm and gneconifer arrangements in Fig. 2C. With the pre-2000 matrices, support and topology are mostly in agreement with the MP analyses. However, with the post-2000 matrices we observe a shift in support from the gnetangiosperms to a clade of Gnetales and conifers. This is illustrated by a split network consensus based on the Rothwell and Stockey (2016) matrix (Fig. 3C), in which Gnetales are linked with conifers, and *Glossopteris*, *Caytonia*, and *Petriellaea* (a Triassic fossil not included in earlier analyses that is now better known vegetatively thanks to work of Bomfleur et al. 2014) are the closest outgroups of angiosperms.

**Figure 3.**
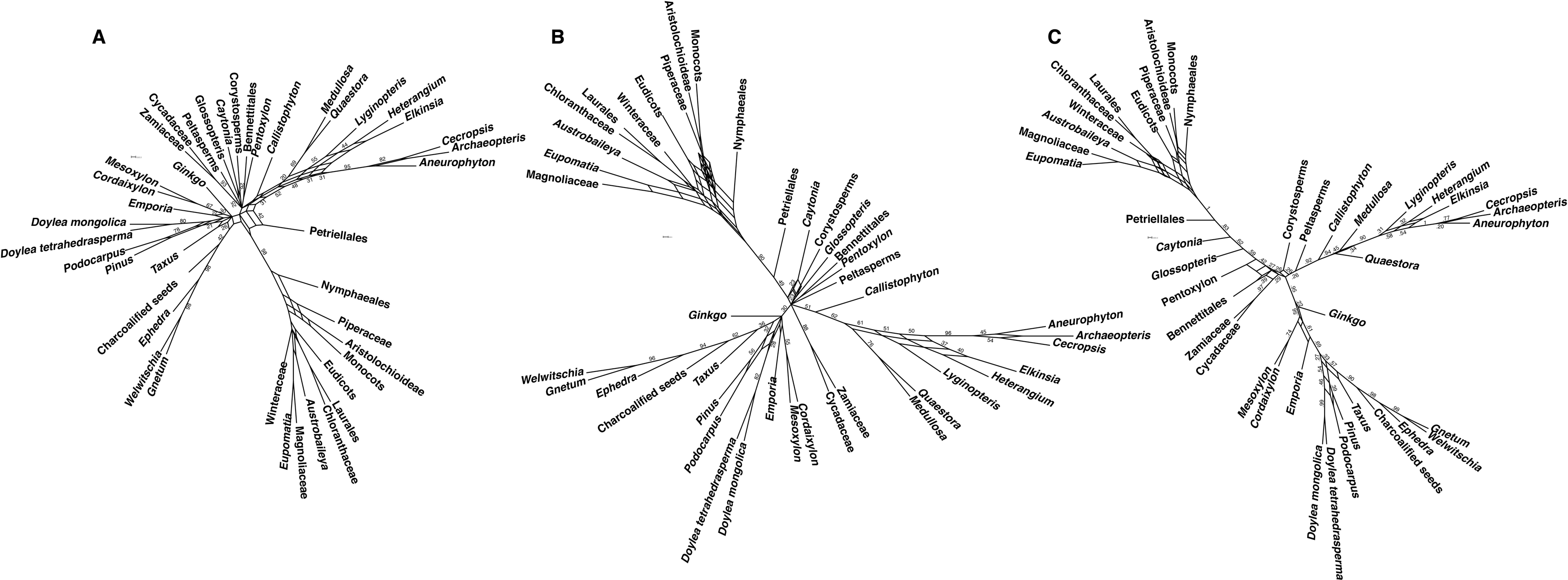
Split network consensus of A, the posterior tree sample of the MP bootstrap analysis, B, the ML bootsrap analysis, and C, the BI analysis of the Rothwell and Stockey (2016) matrix using gamma-distributed rate variation. Only splits with more than 0.15 PP or 15% boostrap are shown, and support is shown only for splits with more than 0.20 PP or 20% bootstrap. The support values for the splits within the angiosperms have been removed for clarity. If the Gnetales-conifer clade is present and supported with all three methods, other relationships (i.e., *Caytonia* in a clade with angiosperms) are only supported in the BI analysis.

Our first test of the hypothesis that the gnetangiosperm topology is the result of long branch attraction consists of long branch extraction (LBE) experiments (Fig. 4A,B). These involved separate removal of the two potential long branch taxa: angiosperms and Gnetales.

**Figure 4.**
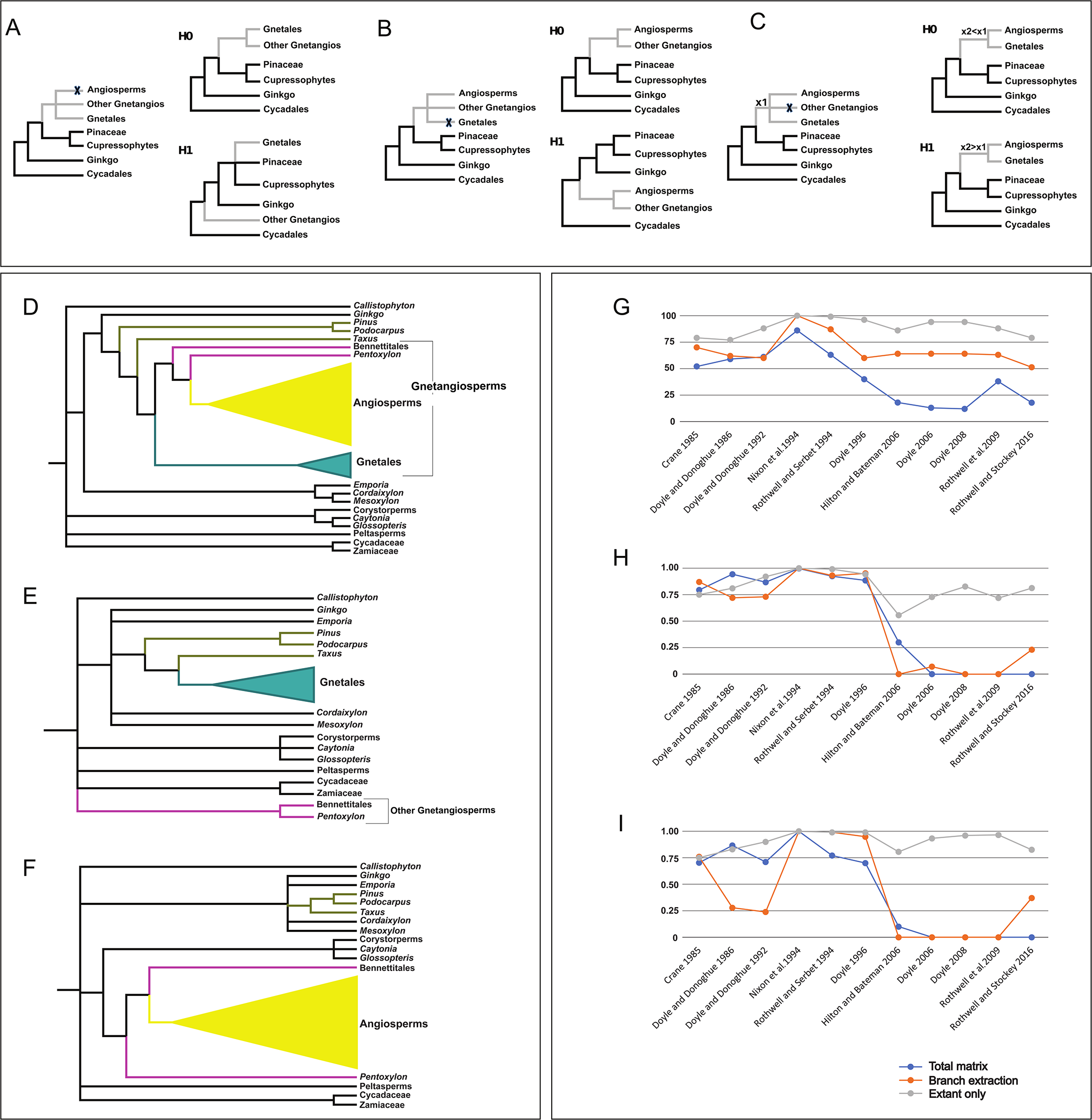
A-C, Scheme of the long branch attraction tests; A and B represent the long branch extraction experiment, C represents the branch elongation experiment. Null hypotheses are in the right upper corner. D-F, Results of the LBE experiment on the Rothwell et al. (2009) matrix. All trees are MP consensus trees. Fossil taxa diverging below the most recent common ancestor of extant seed plants removed for ease of comparison. D, Untrimmed matrix, showing an gnetangiosperm topology and paraphyletic conifers. E, Angiosperm removal matrix, showing Gnetales nested in the conifers and remaining gnetangiosperms removed from the coniferophyte clade. F, Gnetales removal matrix, with monophyletic conifers nested in a large coniferophyte clade. G-I, Results of the BE and EX experiments. G, Results of the MP analyses. H-I, Results of the BI analyses under the Markov k-states (Mk) model with H, equal rates and I, gamma distributed rate variation.

The removal of the angiosperms has different effects on the pre- and post-2000 matrices. With the Crane (1985a) version two matrix analyzed here, a topology with Bennettitales, *Pentoxylon* and the Gnetales diverging after *Lyginopteris* and before the other taxa becomes as parsimonious as the topology with the gnetangiosperms nested among Mesozoic seed ferns that was retrieved with the full matrix. The new tree corresponds to the most parsimonious tree that Crane (1985a) found with his version one matrix, which differed in that Bennettitales and *Pentoxylon* were scored as not having cupules potentially homologous with those of Mesozoic seed ferns. With the Doyle and Donoghue (1986) matrix, Bennettitales, *Pentoxylon*, and Gnetales are nested within coniferophytes. With the Doyle and Donoghue (1992) and Rothwell and Serbet (1994) matrices, the consensus tree is identical to the trimmed consensus derived from the full matrix. With the Nixon et al. (1994) matrix, *Cordaites* and *Ginkgo* are successive outgroups to a conifer plus gnetangiosperm clade, whereas with the full matrix they are equally parsimoniously placed as successive outgroups to the conifers, in a clade that is sister to gnetangiosperms. The inverse happens with the Doyle (1996) matrix, where the position of *Ginkgo* and cordaites is destabilized by the removal of the angiosperms, with these taxa being either successive outgroups to extant and fossil conifers or sister to a clade composed of other former gnetangiosperms, conifers, *Peltaspermum*, and *Autunia.* The position of the Gnetales in a truncated gnetangiosperm clade (i.e., with Bennettitales and *Pentoxylon*) is maintained in all matrices.

With the post-2000 matrices, the effect of removal of the angiosperms is consistent among different matrices. With the Hilton and Bateman (2006) matrix, Gnetales are equally parsimoniously placed within the coniferophytes, within the coniferophytes as sister to the Bennettitales or in an antophyte clade as sister to the conifers. In the Doyle (2006) and Doyle (2008) datasets, the resulting trees see the Gnetales nested within the coniferophytes, with or without Bennettitales. With the Rothwell et al. (2009) matrix (Fig. 4D-F), a topology with a clade of Gnetales and conifers that excludes Bennettitales and *Pentoxylon* becomes most parsimonious (Fig. 4E). With the Rothwell and Stockey (2016) matrix, Gnetales are sister to *Taxus* in a coniferophyte clade that also includes *Doylea*, an Early Cretaceous cone-like structure interpreted as consisting of seed-bearing cupules (Stockey and Rothwell 2009; Rothwell and Stockey 2016).

The removal of the Gnetales has no impact at all on trees based on the Crane (1985a), Doyle and Donoghue (1986), and Doyle and Donoghue (1992) matrices, in which the topology is identical to the trimmed topology of the consensus in the full analysis. With the Nixon et al. (1994) matrix, the removal of the Gnetales results in trees in which coniferophytes form a clade (including *Ginkgo* and *Cordaites*), i.e., eliminating most parsimonious trees in which gnetangiosperms are linked with conifers. With the Rothwell and Serbet (1994) matrix, the removal of Gnetales results in a breakup of the *Caytonia*-*Glossopteris*-corystosperm clade, with the angiosperms still nested within the other gnetangiosperms. With the Doyle (1996) matrix, the only difference lies in the placement of the corystosperms, *Autunia*, and *Peltaspermum*, which are sister to a coniferophyte clade in the analysis without Gnetales.

With the post-2000 matrices, the removal of the Gnetales results in trees in which the remaining gnetangiosperms (which may or may not include *Caytonia*) form a clade outside the coniferophytes (e.g., Fig. 4F). With the Doyle (2006) and Doyle (2008) matrices, a clade including Cycadales, glossopterids, and remaining gnetangiosperms (including *Caytonia*) is sister to a clade of *Callistophyton*, *Peltaspermum*, *Autunia*, and corystosperms plus coniferophytes. The analysis of the Rothwell and Stockey (2016) matrix represents an exception, where the placement of the remaining gnetangiosperms is not affected by the removal of Gnetales. However, the removal of *Doylea* in addition to Gnetales results in a pattern similar to that found with the other post-2000 matrices.

In the branch elongation (BE) experiment, where three fossils commonly associated with angiosperms (Bennettitales, *Pentoxylon*, *Caytonia*) were removed, we observed that MP bootstrap support for the angiosperm plus Gnetales clade increases in all matrices (Fig. 4G). This effect is even stronger in the extant (EX) experiment matrices, in which all fossil taxa were removed, where a split including angiosperms plus Gnetales is strongly supported by the MP bootstrap in all matrices.

Bayesian analysis (BI) of the BE and EX matrices shows a less linear pattern (Fig. 4H, I). In the BE analyses, the signal for the gnetangiosperms decreases with the Doyle and Donoghue (1986, 1992) matrices, reaching less than 0.5 posterior probability (PP) in the analysis with gamma distributed rate variation. With the Nixon et al. (1994), Rothwell and Serbet (1994), and Doyle (1996) matrices, the PP of the gnetangiosperms in the BE matrices is comparable to that from the full matrices. In the post-2000 BE matrices, BI support for the gnetangiosperms is almost null with the Hilton and Bateman (2006) and Doyle (2006) matrices (<0.07 PP) and increases with the Doyle (2008) and Rothwell et al. (2009) matrices analyzed using gamma-rate variation (0.55 and 0.51 respectively) and with the Rothwell and Stockey (2016) matrix (0.23 for the equal-rate analysis, 0.37 for the gamma analysis). The analyses of the EX matrices all show high to moderate support (1-0.75 PP) for the split containing angiosperms plus Gnetales. With the post-2000 matrices, the use of the gamma-distributed model recovers a higher PP for the gnetangiosperms.

The morphospace analyses (Fig. 5) provide a graphic confirmation of the morphological separation of both Gnetales and angiosperms from other seed plants and the perception that Gnetales share competing morphological similarities with both angiosperms and conifers. In the morphospace generated from most of the pre-2000 matrices, Gnetales lie closer to angiosperms (data not shown). With the Doyle (1996) matrix and the post-2000 matrices, the first PCO axis appears to separate angiosperm-like and non-angiosperm-like taxa, whereas the second axis seems to represent a tendency from a seed fern-like towards a conifer-like morphology. Gnetales are always placed closer to the conifers than to the angiosperms (Fig. 5). However, in all cases, Gnetales seem to have higher levels of “angiosperm-like” morphology than do conifers, represented by their rightward placement on the first PCO axis. This position on the first axis is shared by *Doylea* with the Rothwell and Stockey (2016) matrix. Between the analyses of the Doyle (1996) and Doyle (2008) matrices (Fig. 5A, B), there is a modest shift of Gnetales away from angiosperms and towards conifers.

**Figure 5.**
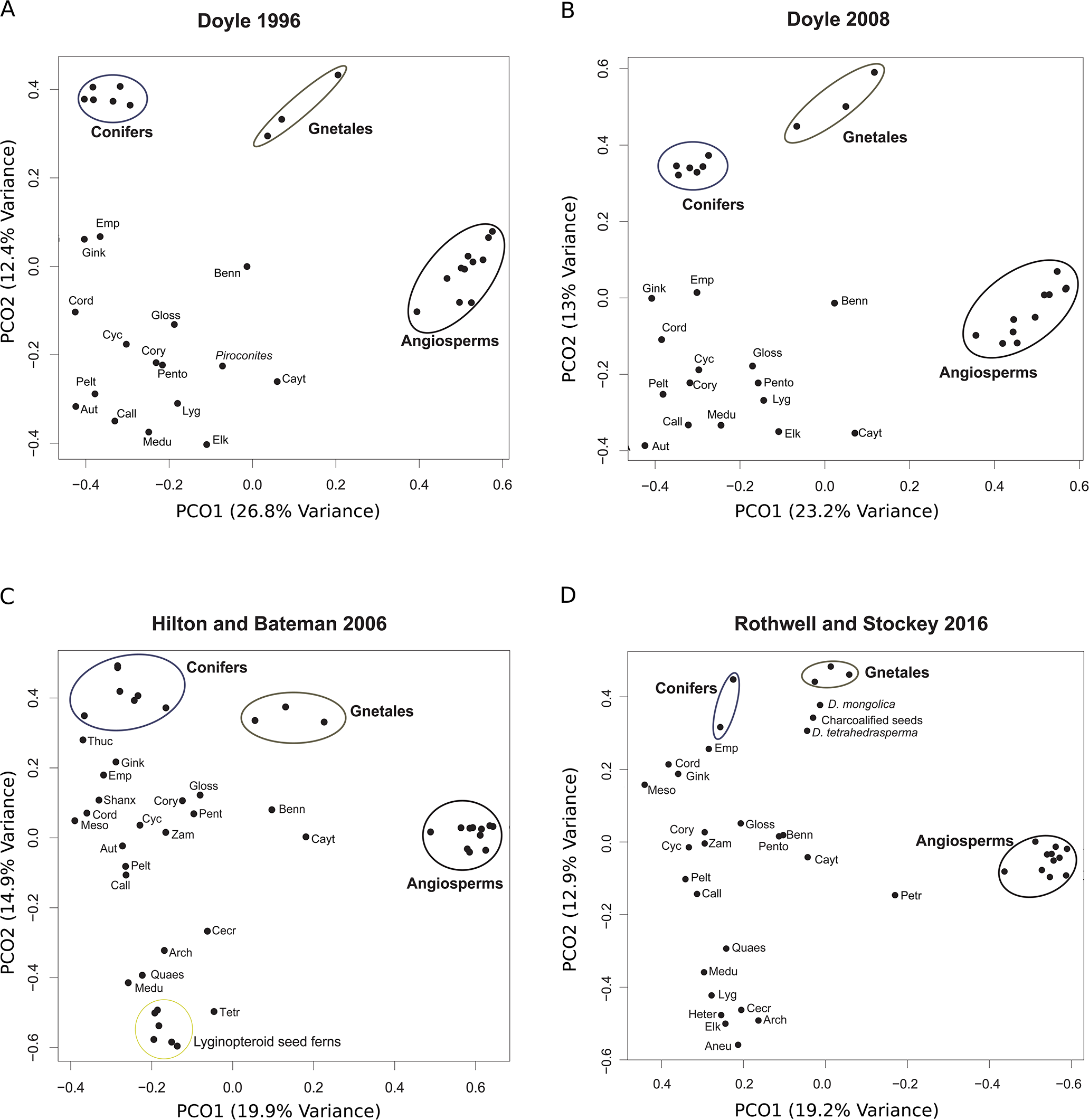
Plot of the first two principal coordinate axes for four of the matrices analyzed. The first PCO axis mainly separates the angiosperms and the other seed plants, while the second PCO axis separates more conifer-like and more fern-like groups. These plots illustrate the effect of the reassessment of gnetalean characters between the two Doyle matrices (A, B), and the similar structure of the data in the Hilton and Bateman (2006) (C) and Rothwell and Stockey (2016) (D) matrices.

## DISCUSSION

The results of our analyses help to resolve some of the main issues regarding the phylogenetic signal for the gnetangiosperm clade in morphological matrices of seed plants. Our meta-analyses of published datasets (Fig. 2) show a two-step trend: first, changes in character sampling and analysis weakened support for the gnetangiosperm hypothesis, and second, the use of model-based methods shifted the balance in favor of a relationship between Gnetales and conifers, bringing the results in line with molecular data. The effect of changes in character analysis is seen in the switch in support between matrices compiled before the main molecular analyses of seed plant phylogeny (pre-2000) and afterwards: i.e., Doyle (2006) and Hilton and Bateman (2006). These two matrices, which both used Doyle (1996) as a starting point but were modified independently, with only limited discussion at later stages of the two projects, and made different choices regarding character coding, taxon sampling, and splitting of higher-level taxa, both show a very similar pattern. Under the MP criterion, a gnetangiosperm topology continued to be more parsimonious, but with reduced support. By contrast, ML and the Bayesian criterion positively favor a grouping of Gnetales and conifers. The matrices descended from Doyle (2006) (i.e., Doyle 2008) and from Hilton and Bateman (2006) (i.e., Rothwell et al. 2009, 2016) exhibit a similar pattern, except that in Doyle (2008) gnetangiosperm and gneconifer trees were equally parsimonious. This phenomenon was already reported by Mathews et al. (2010), who reanalyzed the matrix of Doyle (2008) using BI.

### Critical character reassessment weakened the conflict between morphology and molecules

Examination of the behavior of characters on gnetangiosperm and gneconifer trees illustrates how changes in character analysis made between the studies of Doyle (1996) and Doyle (2006, 2008) increased support for gneconifer trees. Some of these changes were the result of new discoveries concerning the morphology of Gnetales and other taxa, others of critical reassessment of previous character definitions aimed at reducing bias in favor of the gnetangiosperm hypothesis. The shift of Gnetales away from angiosperms and towards conifers observed in the morphospace analyses based on the datasets of Doyle (1996) and Doyle (2008) (Fig. 5A, B) is presumably the result of these changes. Especially modifications of the latter sort illustrate general problems of analysis and definition of morphological characters, which can be far more difficult than is usually acknowledged. Because potentially homologous structures in different taxa differ to various degrees, there is often a tension between use of overly lax criteria for definition of states at the stage of primary homology assessment, which may mistake homoplasy for homology, and overly strict criteria, which may overlook real synapomorphies. Other problems can be caused by inclusion of distinct characters that are correlated for functional or developmental reasons and therefore overweight single transformations, or by decisions on whether to treat presence and absence of a structure and different forms of the structure as states of the same character or as separate characters, both of which can lead to artifacts.

Most changes of the first sort involved previously overlooked conifer-like features of Gnetales. For example, Doyle (2006) added a character for presence of a torus in the pit membranes of xylem elements in conifers and Gnetales, based on observations on Gnetales by Carlquist (1996) and studies of conifers by Bauch et al. (1972). Doyle (2006) also rescored Gnetales as having a tiered proembryo, as in conifers; two tiers of cells were illustrated by Martens (1971) and called “étages,” and by Singh (1978). This similarity may have been overlooked because of other differences related to elimination of a free-nuclear phase in the embryogenesis of Gnetales (Doyle 2006). Both characters undergo one less step on gneconifer trees than on most gnetangiosperm trees (exceptions are some trees with major rearrangements elsewhere in seed plants). In male “flowers” of *Ephedra* and *Welwitschia*, microsynangia are borne in two lateral groups, which Doyle (1996) interpreted as reduced pinnate sporophylls. Because Bennettitales, *Caytonia*, and many “seed fern” outgroups have pinnately organized microsporophylls, this character favored a gnetangiosperm tree by one step. However, developmental studies by Mundry and Stützel (2004) indicated that the two lateral structures are more likely branches (strobili) bearing three or four simple sporophylls. Based on these observations, Doyle (2008) rescored microsporophylls in Gnetales as simple and one veined, as in conifers, and as a result the character favored the gneconifer topology by one or two steps.

Doyle (2006) also made changes based on improved data on a character expressing the position of the ovule or ovules on the sporophylls or “cupules” that bear them, which is not directly relevant to Gnetales but potentially useful for identification of gnetangiosperm outgroups. Ovules are on the abaxial surface of the sporophyll/cupule in corystosperms (Axsmith et al. 2000; Klavins et al. 2002), rather than on the adaxial surface in glossopterids (Taylor and Taylor 1992), probably *Caytonia*, and angiosperms (if the outer integument is a modified leaf or cupule: Doyle 2006, 2008; Kelley and Gasser 2009). Ovules are also adaxial in the cupules of *Petriellaea* (Taylor et al. 1994; Bomfleur et al. 2014), which was included in the analysis of Rothwell and Stockey (2016).

Other changes were the result of doubts concerning the homology of characters that supported the gnetangiosperm hypothesis, along lines suggested by Donoghue and Doyle (2000). For example, in the apical meristem character, Doyle (1996) contrasted the presence of a tunica (an outer layer that maintains its integrity by undergoing only anticlinal cell divisions, i.e., perpendicular to the surface) of Gnetales, angiosperms, and Araucariaceae, vs. its absence in cycads, *Ginkgo*, and other conifers. This character undergoes two steps when Gnetales are linked with angiosperms (the state in fossils is unknown), three when Gnetales are linked with conifers. However, the tunica consists of one layer of cells in Gnetales, but two layers in angiosperms, suggesting that it may not be homologous in the two groups. To reduce bias in favor of homology of these two conditions, Doyle (2006) split presence of a tunica into two states. The resulting three state character undergoes three steps with Gnetales in both positions. Redefinition of the megaspore membrane character involved a shift in the limit between states, from thick vs. reduced (thin or absent) to present vs. absent; the megaspore membrane is thin in Gnetales, but absent in angiosperms, *Caytonia*, and probably Bennettitales. In compressions of bennettitalean seeds prepared by oxidative maceration, Harris (1954) observed no megaspore membrane, but Wieland (1916) and Stockey and Rothwell (2003) reported a thin layer around the megagametophyte in permineralized seeds. However, as noted by Harris (1954), there is no evidence that this layer is a true megaspore membrane (i.e., consisting of exinous material). These changes in character definition do involve a subjective element and were doubtless influenced by knowledge of the molecular evidence for a relationship of Gnetales and conifers, but the new definitions represent a shift toward greater caution in evaluating the potential homology of similar but not identical structures.

The trends seen in Fig. 2 show that recognition of previously overlooked similarities between Gnetales and conifers and reconsideration of potentially convergent characters between angiosperms and Gnetales succeeded in strengthening a morphological signal associating Gnetales with conifers. This result clearly contradicts the view that morphology and molecules are in strong conflict with each other (Bateman et al. 2006; Rothwell et al. 2009) and validates arguments along these same lines that were advanced by Doyle (2006, 2008) on a parsimony basis. Indeed, in all post-2000 matrices a topology with Gnetales linked with conifers requires the addition of only a few steps to the length of gnetangiosperm trees: e.g., four in the case of Hilton and Bateman (2006) and one in Doyle (2006), and in Doyle (2008) both topologies became equally parsimonious. A tendency to focus on the MP consensus tree and lack of exploration of almost equally parsimonious alternatives may have tended to inflate the perceived conflict between molecules and morphology. Among analyses since 1994, bootstrap and/or decay values were reported by Doyle (1996, 2006, 2008), Hilton and Bateman (2006), and Rothwell and Stockey (2016), but not by Nixon et al. (1994), Rothwell and Serbet (1994), and Rothwell et al. (2009). Our analyses show that the signal retrieved using MP is more correctly characterized as profoundly ambiguous.

### Contribution of model-based methods

By contrast, maximum likelihood and especially Bayesian analyses of all post-2000 matrices converge on a similar result, unambiguously favoring placement of Gnetales in a coniferophyte clade that includes Ginkgoales, cordaites, and extant and extinct conifers. Stronger support is obtained in BI analyses in which gamma rate variation among sites is implemented in the model. With ML the difference in relative support for the two hypotheses appears smaller, but a gneconifer arrangement is consistently favored with all datasets. These results of model-based analyses of post 2000 morphological matrices have interesting implications regarding stem relatives of the angiosperms. Indeed, most post-2000 matrices are broadly congruent in attaching *Pentoxylon*, glossopterids, Bennettitales, and *Caytonia* to the stem lineage of the angiosperms. To these the analysis of Rothwell and Stockey (2016) adds the Triassic genus *Petriellaea* (Taylor et al. 1994; Bomfleur et al. 2014) (Fig. 3), which has simple reticulate laminar venation, as in *Caytonia*, and cupules containing adaxial ovules. This may be consistent with the view that these fossils shed light on evolution of the complex reticulate venation and bitegmic ovules of angiosperms (Doyle 2006, 2008).

A cautionary note on the results of our Bayesian analyses is necessary. The differences between bootstrap support values in the MP and ML analyses and posterior probabilities in the BI analyses could be due to the very different nature of these support metrics. It has been shown that the relationship between character support and increase in PP is far from linear, and PP can easily sway results toward a hypothesis that is supported by only a few characters (Zander 2004). The strong PP support for groupings (like *Caytonia* or *Petriellea* plus angiosperms) that receive weak or non-existent support on a character basis (MP and ML bootstrap, Fig. 3A, B) could indicate either the ability of Bayesian inference to pick up a significant signal in an otherwise noisy background or the possibility that this method can be led astray by a few potentially unimportant characters.

### The conflict between morphology and molecules is partially due to long branch attraction

Our results also add new empirical evidence on debates concerning the strengths and weaknesses of morphological data in reconstructing phylogenetic relationships, the phylogenetic importance of fossils, and the best methods to analyze morphological data (Wright and Hillis 2014; O’Reilly et al. 2016; Puttick et al. 2017a). A well-known cause of phylogenetic conflict is the presence of long branches in the tree, which can lead to LBA phenomena (Felsenstein 1978; Bergsten 2005). Analyses based on simulated matrices and real data have repeatedly shown that probabilistic, model-based approaches are more robust to LBA than parsimony (Swofford et al. 2001; Brinkmann et al. 2005; and references therein). Long branch attraction is most commonly discussed as a confounding factor in molecular studies, as in the case of Gnetales-basal trees found with molecular data (Sanderson et al. 2000; Magallón and Sanderson 2002; Burleigh and Mathews 2007), but here it is morphology that is potentially affected: the BI trees show that both angiosperms and Gnetales are situated on very long morphological branches, especially in the post 2000 matrices.

After following suggestions by Bergsten (2005) and other methodologies (Rota Stabelli et al. 2011), we conclude that LBA is responsible at least in part for the continuing support for the gnetangiosperm clade in MP analyses of the post-2000 matrices. First, BI recovers a gneconifer topology with higher probability than a topology with Gnetales linked with angiosperms, thus favoring a topology that separates the long branches over a topology that unites them. Second, more complex and better-fitting models recover a higher posterior probability for the topology in which angiosperms and Gnetales are separated (Fig. 2C). Third, removing Gnetales or angiosperms results in a rearrangement of the MP topologies in which the other long branch “flies away” from its original position. Fourth, support for Gnetales plus angiosperms increases with decreased sampling of fossil taxa on the branch leading to the angiosperms, and still more with the removal of all fossils (Fig. 4G-I). It has been suggested that molecular analyses may be incorrect about the relationship of angiosperms and Gnetales because they ignore the great diversity of extinct seed plant taxa (e.g., Rothwell et al. 2009). This reasoning seems to assume that addition of fossils would strengthen the gnetangiosperm hypothesis, but in fact our results indicate that the opposite is true.

To our knowledge, this represents the first reported case of LBA in a morphological analysis that is supported by multiple tests (Bergsten 2005), with much stronger support than in previously reported cases (Lockhart and Cameron 2001; Wiens and Hollingsworth 2000). These analyses also support the view that model-based methods can overcome the shortcomings of parsimony in such cases. It is also noteworthy that the impact of LBA can be easily visualized with the principal coordinates analysis (Fig. 5), where the presumed close relationship between Gnetales and conifers and the convergence of Gnetales with the angiosperms are effectively congruent with the positions of the three taxa in the plot of the first two PCO axes. This tool could represent an interesting option for exploring the structure of the data in future phylogenetic analyses.

Less intensive examination of our results suggests that there are fewer conflicts between relationships obtained with parsimony and model-based approaches in other parts of the seed plant tree, suggesting that MP is not necessarily misleading when long branch effects are lacking. Even when morphological parsimony analyses vary in the arrangement of extant seed plant lines, they are more consistent about relationships below the crown group, with “progymnosperms,” hydrasperman “seed ferns,” *Lyginopteris*, and medullosans diverging successively below the crown group, and our model-based trees show similar relationships. Another consistent result is the association of traditional coniferophyte groups, namely ginkgos, cordaites, and conifers, setting aside whether this clade also includes Gnetales or (in some morphological analyses) gnetangiosperms. Relationships among cycads and Permian and Mesozoic “seed ferns” (peltasperms, corystosperms, glossopterids, *Caytonia*) are more variable among parsimony analyses, possibly because of the smaller proportion of preserved characters in the fossils and/or the low number of changes on short internal branches between these lines. Assuming that molecular and model-based morphological results are correct, these considerations suggest that parsimony may perform well when branch lengths are moderate, and it would be unwarranted to reject results out of hand because they are based on parsimony.

The conclusion that similarities between angiosperms and Gnetales are the result of convergence should not be difficult to accept, because many aspects of the morphology of Gnetales can be explained in terms of a Paleozoic conifer prototype (which had female branch systems with secondary short shoots bearing sterile and fertile appendages; cf. Rothwell and Stockey 2013). However, removal of Gnetales from the former gnetangiosperm clade introduces new problems, notably by implying that similarities in seed morphology and anatomy in Gnetales and Bennettitales emphasized by Friis et al. (2009) are also convergences. Some of these similarities have been questioned or reduced by subsequent studies of Bennettitales (Rothwell et al. 2009; Doyle 2012; Rothwell and Stockey 2013; Pott 2016), but others remain. These similarities could be homologous if Bennettitales and Gnetales formed a clade within conifers, but it is much less plausible to interpret Bennettitales as modified conifers, considering their cycad-like leaf morphology, wood anatomical features, and pinnate microsporophylls.

## CONCLUSIONS

The main lesson of our analyses may be that, contrary to previous impressions, morphological data do not present a strong conflict with the results of molecular analyses regarding the position of angiosperms and Gnetales. This strongly suggests that morphology carries a phylogenetic signal that is consistent with molecular data, and may therefore be useful in reconstructing other aspects of the phylogenetic history of the seed plants, most notably the position of fossils relative to living taxa. The supposed conflict between the two sorts of data on the major aspect of the phylogeny of seed plants emphasized here seems to be due to a combination of difficult problems in character analysis and limitations of phylogenetic methods. Since data from the fossil record are particularly important for resolving the evolutionary history of seed plants, because of the wide gaps that separate extant groups and the potential biases in analysis of such sparsely sampled taxa (Burleigh and Mathews 2007; Mathews 2009; Rothwell et al. 2009; Magallón et al. 2013), our results give new hope for the possibility of integrating fossils and molecules in a coherent way. This is even more important in light of new fossil discoveries (e.g., Rothwell and Stockey 2013, 2016), some of which show similarities to fossils previously associated with angiosperms (e.g., the Triassic *Petriellaea* plant, which shares leaf and cupule features with *Caytonia*: Bomfleur et al. 2014).

The absence of deep convergence problems also opens the possibility of combining morphological and molecular datasets in a total-evidence analysis. Such an approach has been rarely employed in datasets with fossil and extant plants (Magallón 2010), but it has proven to be useful in resolving some controversial relationships (i.e, in the Cycadales: Coiro and Pott 2017). However, especially with the recent expansion in the amount of available molecular data, both marker selection and taxon choice would have to be carefully considered to set up a successful analysis. It is possible that the ever-increasing amount of sequence data used to infer phylogenetic relationships could swamp the signal present in the many fewer morphological characters, in which case the result would not differ from that found with use of a molecular backbone constraint tree.

An important general message that emerges from our study is the importance of including an exploration of the signal in all phylogenetic analyses involving morphology. The overreliance on single consensus trees, as discussed in Brown et al. (2017) and Puttick et al. (2017b), has been a major driver of the perceived conflict in seed plant phylogeny; another factor has been the lack of support statistics in many studies. Among methods of signal dissection, consensus networks and distance-based neighbor-nets (even if these suffer from the general shortcomings associated with distance-based methods) present promising avenues for the exploration of morphological datasets (Bryant and Moulton 2004) and have proven their power in understanding the history of different groups of fossil and extant taxa at different taxonomic scales (Denk and Grimm 2009; Bomfleur et al. 2017; Grimm 2017).

Although most phylogenetic analyses based on morphology are still conducted in a parsimony framework, some authors have already underlined the potential of model-based approaches in this field (Lee and Worthy 2012; Lee et al. 2014). Our analyses show that BI yields more robust results under different taxon sampling strategies, and although parsimony and BI usually give congruent results, BI appears to be effective in correcting errors of parsimony analyses caused by long branch effects. Our study converges with previous work indicating that the use of model-based techniques could allow the successful integration of taxa with a high proportion of missing data (Wiens 2005; Wiens and Tiu 2012), which is a prime consideration when dealing with the paleobotanical record.

## SUPPLEMENTARY MATERIAL

The supplementary material is available as an online appendix.

## ACKNOWLEDGMENTS

MC acknowledges H. Peter Linder for his fundamental support to this work, and for important comments on this manuscript. We would like to thank Richard Bateman and Gar Rothwell for making their matrices available, and Omar Rota-Stabelli for useful discussions of long branch attraction. Guido Grimm is thanked for thorough comments on the manuscript and a detailed discussion of the use of phylogenetic networks in the analysis of morphological data. Tanja Stadler, Susanne Renner, Elisabeth Truernit, Gavin George, Frank Anderson, Erika Edwards, and two anonymous reviewers are gratefully acknowledged for comments on a previous version of this manuscript. Guy Atchison, Yanis Bouchenak-Khelladi, Merten Ehmig, Kevin Boyce, Catarina Rydin, and an anonymous reviewer are thanked for useful comments on the present version.

